# Response to thermal and infection stresses in an American vector of visceral leishmaniasis

**DOI:** 10.1101/2020.11.23.373100

**Authors:** Kelsilandia A. Martins, Caroline S. Morais, Susan J. Broughton, Claudio R. Lazzari, Paul A. Bates, Marcos H. Pereira, Rod J. Dillon

## Abstract

The phlebotomine sand fly *Lutzomyia longipalpis* is the primary insect vector of visceral leishmaniasis in the Americas. For ectothermic organisms such as sand flies, the ambient temperature is a critical factor influencing all aspects of their life. However, the impact of temperature has been ignored in previous investigations of stress-induced responses by the vector, such as taking a blood meal or during *Leishmania* infection. Therefore, this study explored the interaction of *Lu. longipalpis* with temperature by evaluating sand fly behaviour across a thermal gradient after sugar or blood-feeding, and infection with *Leishmania mexicana.* Thermographic recordings of sand fly females fed on mice were analysed, and the gene expression of heat shock proteins HSP70 and HSP90(83) was evaluated when insects were exposed to extreme temperatures or infected. The results showed that 72h after blood ingestion females of *Lu. longipalpis* became less active and preferred relatively low temperatures. However, at later stages of blood digestion females increased their activity and remained at higher temperatures prior to taking a second blood meal; this behaviour seems to be correlated with the evolution of their oocysts and voracity for a second blood meal. No changes in the temperature preferences of female sand flies were recorded in the presence of a gut infection by *Le. mexicana,* indicating that this parasite has not triggered behavioural immunity in *Lu. longipalpis*. Real-time imaging showed that the body temperature of female flies feeding on mice increased to the same temperature as the host within a few seconds after landing. The body temperature of females remained around 35 ± 0.5 °C until the end of blood-feeding, revealing a lack of thermoregulatory behaviour. Analysis of expression of heat shock proteins revealed insects increased expression of HSP90(83) when exposed to higher temperatures, such as during blood feeding. Our findings suggest that *Lu. longipalpis* interacts with the environmental temperature by using its behaviour to avoid temperature-related physiological damage during the gonotrophic cycle. However, the expression of certain heat shock proteins might be triggered to mitigate against thermal stress in situations where a behavioural response is not the best option.

## 1. Introduction

In Latin America, the sand fly *Lutzomyia longipalpis* is the primary insect vector of *Leishmania infantum*, the causative agent of visceral leishmaniasis in this region of the world (Lainson and Rangel, 2005). The transmission of parasites usually happens when infected females take another blood-meal and inject the metacyclic promastigote form of *Leishmania* into the host. Along with many other species of sand flies, *Lu. longipalpis* is found in tropical areas and lab investigations suggest an optimum development for this species between 20-28°C (Guzmán and Tesh, 2000).

The ambient temperature is an abiotic factor that can directly influence the fitness of insects, affecting their development, reproduction and survival (Bale et al., 2002). For hematophagous insects as sand flies, additional changes in their body temperature are expected considering their blood-feeding on various homoeothermic vertebrates. For example, the body temperature of mammals can vary in a range of 30-38°C, whilst for birds it is 37-43°C (Clarke, 2007). Moreover, blood-sucking insects have evolved to ingest a large volume of blood within a short period to escape host defences. Therefore, blood-feeding results not only in a dramatic increase in their body size but also potentially temperature in a few minutes (Lahondère and Lazzari, 2012). A rapid change in the body temperature of insects can impair protein synthesis, metamorphosis cessation, cell division, and hormone secretion among insects. Therefore, many of them have developed several strategies to avoid the risk of thermal stress (Heinrich, 1993; Okasha, 1968).

Insects usually can adapt to temperatures that differ from their own needs by adopting two typical strategies of thermoregulation mechanisms: regulation of body temperature by their behaviour or by making biochemical changes according to the external thermal environment (Matthew and Blanford, 2003). From the molecular point of view, the participation of proteins, known as Heat Shock Proteins (HSPs), has been described in several organisms during stress situations including those caused by extreme temperatures (Lu and Wan, 2010). These proteins assist the thermal defence system by acting as chaperones, having an important role in protecting and maintaining essential cellular functions (Marrof et al., 2005). According to their molecular weight, HSPs are classified into different families (HSP100, HSP90, HSP70, HSP60 and small HSPs) (Saidi et al.,2009). Several of these families have been described in situations when insects were exposed to extreme temperatures inducing a thermal tolerance response (Benoit et al., 2009).

The regulation of body temperature is also a well-described behavioural immunity response among the defence systems of insects against pathogens (Roode and Lefèvre, 2012). When infected by specific pathogens some insects can manipulate their temperatures by temperature selection in order to impair the parasite, moving to colder or warmer temperatures that differ to their natural range (Hinestroza, 2016; Truitt, 2019). On the other hand, certain parasites are also capable of altering the thermal behaviour of their hosts to improve their virulence and suitability (Matthew and Blanford, 2003).

Recently the impact of temperatures on insects started to attract more attention in those with vectorial capacity, especially because modifications in the epidemiology of vector-borne diseases have been strongly associated with climate changes in recent years (Lin et al., 2019; Liu et al., 2019; Hertig, 2019). Climate conditions are considered as one of the most critical factors influencing diseases such as leishmaniasis, especially concerning the sand fly vectors (Chalghaf et al., 2018; Oliveira et al., 2018). Vectors such as *Lu. longipalpis* tend to be more active in warm environments, which would favour the transmission of the parasite (Rivas et al., 2014). In addition, *Leishmania* parasites are also influenced by the temperature of insect, some species such as *Le. infantum* and *Le. braziliensis* can develop better in *Lu. longipalpis* when maintained at higher temperatures (26°C) (Hlavacova et al., 2013). Conversely, the longevity of this sand fly is increased at cooler temperatures (15°C) (Guzmán and Tesh, 2000). However, all these findings upon parasites or sand flies have been studied with fixed temperatures, disregarding the choices of insects as would occur in nature.

Considering the influence that temperatures can play on behaviour, physiology and immunology of insects, we investigated some of the unknown aspects of the relationship of *Lu. longipalpis* with temperature. This interaction was studied by observing the behaviour of this species during blood-feeding on the host, over its gonotrophic cycle and infection by *Le. mexicana*, and by evaluation of heat shock protein expression in relation related to thermal stress.

## 2. Materials and methods

### 2.1. Sand Flies

Brazilian sand flies (originally from Jacobina and Teresina) were maintained in facilities at Lancaster University (UK) and Instituto de Ciências Biológicas/UFMG (Brazil) using standard temperature conditions (24 ± 2°C), artificial photoperiod, humidity < 70% and providing sugar *ad libitum* to adults. All insects were reared based on modified methodology from Modi and Tesh (1983).

### 2.2. Infections with *Leishmania* mexicana

*Leishmania* infections were performed artificially via a Hemotek membrane feeder (Discovery workshops, UK) at 37°C using sheep blood stored in Alsever’s anticoagulant (Cat. No. SB068, TCS Biosciences, UK). Cultured amastigotes of *Leishmania mexicana* (Reference strain -MNYC/BZ/62/M379) were used for sand fly infections following the same procedures as detailed by Moraes et al. (2018).

### 2.3. Thermal behaviour of *Lu. longipalpis*: locomotor activity in a thermal gradient

Experiments evaluating the temperature preferences of sand flies were performed using a thermal gradient constructed with an ‘L’ shaped aluminium block. The block was connected between a hot plate (Stuart Scientific UK) and a cooling water bath using a thermostatic circulator (LKB, Sweden) to generate a temperature gradient **(Figure 1A).** Before the beginning of experiments, the water bath and the heating plate were switched-on, and after approximately 2 hours a temperature gradient of 18 °C – 30 °C (±1 °C) in the aluminium block was established. This range of temperatures was chosen to simulate the effect of seasonal and environmental variations on the behaviour of *Lu. longipalpis*. The apparatus was calibrated to create a linear gradient of temperatures divided into six stable zones, as shown in **Figure 1B**. At the start of each experiment, the temperature in the centre of each zone was measured using a calibrated k-type thermocouple (Digi Sense, UK) to verify the stability of the gradient. Cold-light lamps on both sides of the apparatus were used to ensure uniform illumination across the device. During experiments sand flies were placed into a white aluminium tray (270 × 180 × 5mm) with a Perspex cover of the same dimensions. The tray was divided into 4 cm wide arenas, and a ruler placed in one of the edges was used to verify the limit of each zone. Small holes were drilled on top of the Perspex cover to allow the release or capture of the insects during each trial, and they were covered with transparent tape to avoid interference with video tracking and escape of insects.

**Figure 1.**
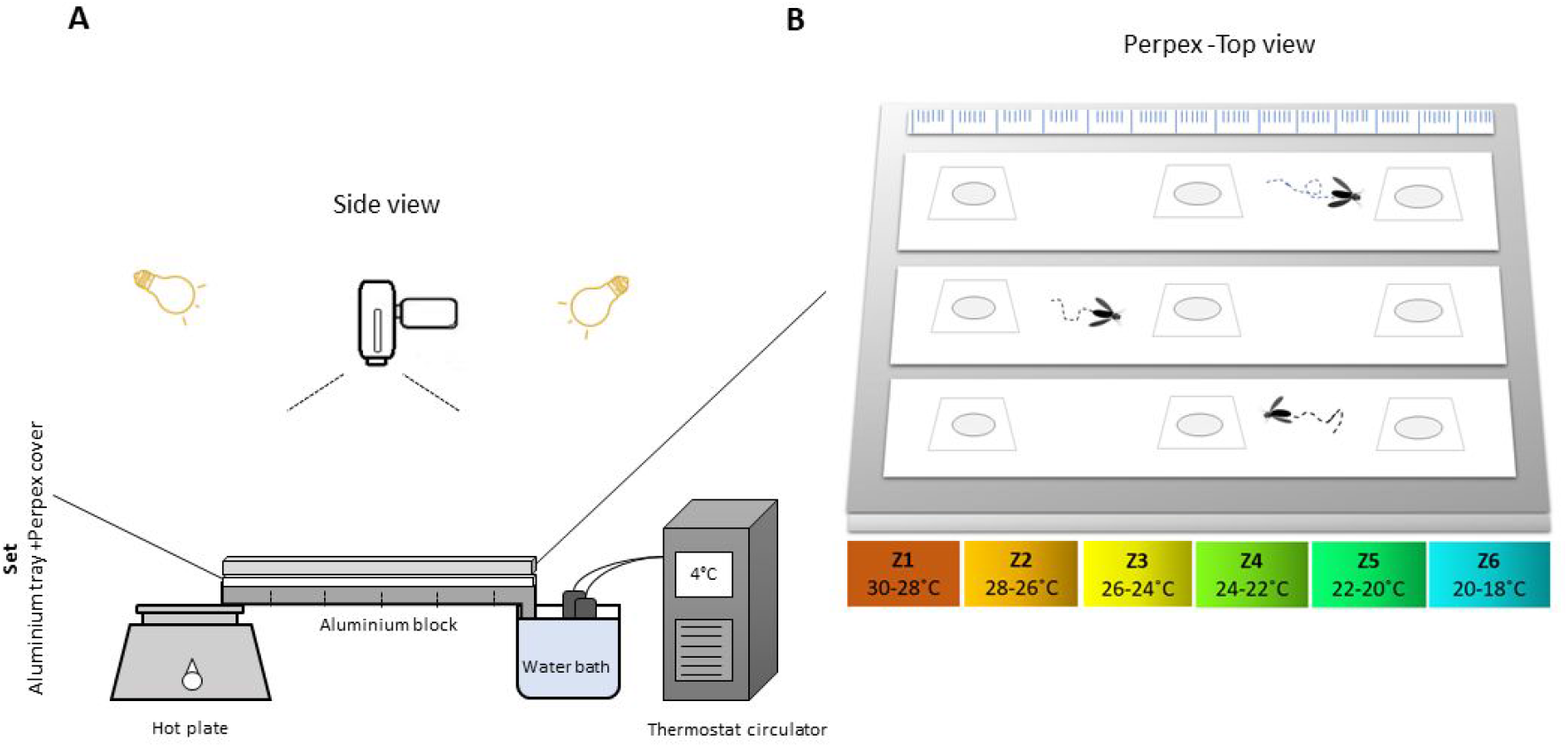
Illustration of the apparatus used during experiments into thermal preferences. **A** The side view of the apparatus used during trials with the description of each component. **B** The top view of arenas and zones of temperature generated in the aluminium tray.

### 2.4. Analysis by video-tracking

All videos were recorded using a video camera (Sony DCR SR8E) and analysed by the EthoVision software animal tracking software (Noldus Information Technology, Wageningen, The Netherlands). Each experiment was recorded for ≈40 minutes, and videos were analysed by the software when an insect was detected in the arena, then set to run for 30 minutes from that point. The first and last 4 minutes of each video were not used in tracking due to time required to stabilise temperatures on the aluminium tray and interferences caused by the operator setting the camera to record or stop. The tracking method used in the analysis recognises the differences in contrast and motion between a static image of the tracks and the video input. The parameters selected for the software were analysed to detect the locomotor behaviour of insects on thermal gradient (duration of time spent in each zone) and activity (mean velocity, distance travelled).

### 2.5. Experimental procedure

Adults were aspirated individually per arena into the middle hole of the plate, then after covering the hole with the tape, the set was transferred to the top of the aluminium block. All experiments were recorded during the same period of the day (≈13:00-18:00) avoiding human presence inside the recording room. Nitrile gloves were used during handling of all equipment to minimise the possibility of contamination by human-derived odours, the room was windowless, and the tray was wiped with ethanol 70% after each trial. When working with infected females, a danger alert tape was placed around the plate and the set was transferred to a freezer at −20 °C for at least 5 minutes, before further dissections of females to confirm the presence of promastigotes. The locomotor behaviour was evaluated using females 3 or 6 days post-ingestion of blood (PBI), and females 3 or 6 days post-infection with *Leishmania* (PIL). Three days after infection the presence of *Le. mexicana* promastigotes in the midgut of females is expected and by 6 days it is possible to find infective forms located in the cardia region of females (Gossage et al., 2003). The behaviour on thermal gradients was also evaluated in sugar-fed (SF) males and females 4 or 7 days old. Before initiating experiments observing these groups, 4 day old females and males (SF) at room temperature (20 °C) were tested with the apparatus turned-off to observe if *Lu. longipalpis* could have a natural preference for a specific section of the arena.

### 2.6. Effect of different temperatures on *Lu. longipalpis* and *Le. mexicana*

We observed the oviposition percentage and survival of females after oviposition in groups kept at 20 °C, 24 °C and 30 °C. We followed these females for six days PBI to describe the effect of temperatures on the behaviour of gravid females . Some females PIL were also dissected to observe the evolution of parasites at these temperatures. Oviposition assays were performed with females maintained inside clear plastic pots (4.6 cm × 4.3 cm) filled with 1 cm of moist Plaster of Paris on the bottom. The top part of the pot was covered with netting fabric with a 1cm diameter hole, plugged with cotton wool to allow the entrance of insects. As soon as each female was aspirated individually into the pot, a dry piece of cotton wool was inserted into the hole to avoid escapes, and a second piece of cotton wool soaked in a sugar solution was placed on top of the fabric. Each group of pots was stored in a plastic box with moistened tissue paper on the bottom and then transferred to the planned temperatures in incubators.

### 2.7. Thermography

The temperature of the *Lu. longipalpis* females was also evaluated during feeding on a hairless mouse using a thermographic camera (PYROVIEW 380L compact, DIAS infrared GmbH, Germany; spectral band: 8-14 mm, without a 2D cooling detector, 384 288 pixels) equipped with a macro lens. The thermal data were evaluated by RTV Analyzer RT software (IMPAC Systems GmbH, Germany), and the body temperature of sand flies was measured frame by frame from the recordings (n=10) (Lahondère and Lazzari, 2012). All hairless mice used in this experiment were kept with free access to water and food at facilities of the Department of Parasitology, Universidade Federal de Minas Gerais-UFMG, Brazil. In each experiment, the mouse was anaesthetised via intraperitoneal injection with 150 mg/kg ketamine (Cristalia, Brazil) and 10 mg/kg xylazine (Bayer, Brazil) in a 1:1 ratio. Mice were kept at a constant temperature on a thermal blanket (Fine Science Tools, Vancouver, Canada) during procedures. The experiments were approved by the Ethical Committee on Animal Use (CEUA/UFMG-Universidade Federal de Minas Gerais-Brazil) under the Protocol n°. 316 / 2015.

### 2.8. Gene expression of Heat Shock Proteins in *Lu. longipalpis*

#### 2.8.1. Primers

Expression of genes potentially stimulated by temperature-stress in *Lu. longipalpis* was evaluated using heat shock protein sequences from families HSP 70 and 90(83). The primers used for conventional PCR were designed using Primer3 software (http://primer3.sourceforge.net/). Oligonucleotides were synthesised based on the homology between the published sequences of *Lu. longipalpis* and the Heat-shock Proteins HSP90(83) from *Drosophila melanogaster* (GenBank: NM-001274433.1) and HSP70 from *Culex quinquefasciatus* (GenBank: XM_001861403.1), evaluated by BLAST. The primers used in the real-time PCR reactions were aligned using the Clustal Omega program (http://www.ebi.ac.uk/Tools/msa/clustalo/) and designed by the NCBI software (http://www.ncbi.nlm.nih.gov/tools/primer-blast/). The efficiency of the primers was tested before reactions by serial dilution of the cDNA using the slope value of the curve under a model linear regression. Efficiencies between 90% and 110% were considered acceptable (Radonic et al., 2004).

#### 2.8.2. PCR protocol and experimental groups

The total RNA was extracted using Trizol reagent (Invitrogen) in pools containing ten insects per sample. After the extraction, the RNA samples were treated with a Turbo DNA-free kit (Life Technologies), following the manufacturer’s instructions. All samples were quantified by spectrophotometer (A260/280nm) in the NanoDrop 2000 UV Spectrophotometer apparatus (ThermoScientific) and stored at −80 °C until use. For the preparation of the cDNA, the RNA was eluted in Milli-Q water and using 0.5 μg of random hexamer primers (Promega) and the M-MLV reverse transcriptase system (Promega) in a final volume of 20 μL. For amplification of the material conventional PCR reactions were performed under the following conditions: 35 cycles (94°C / 30 sec, 60°C / 30 sec, 72°C / 45sec) with 1 μl of cDNA, 150 nM of each primer, 200 μM dNTPs and 1 U of Taq DNA polymerase in a final volume of 20 μL. PCR product analysis was performed by agarose gel electrophoresis at 2.0%, stained with GelRed at 1: 400 (Biotium) and visualised under ultraviolet light. For each evaluated gene, a negative control was made using sterile water instead of cDNA.

#### 2.8.3. Evaluation of gene expression

After synthesis of the cDNA, real-time PCR reactions were performed with the Power SYBR Green PCR MasterMix Kit (Applied Biosystems) following the manufacturer’s recommendations. Each reaction was performed using 12μl of the Power SYBR Green PCR Master Mix (Applied Biosystems), 10ng of cDNA and 300nM of each of the primers. The PCR primers and amplicon sizes were: HSP70-Lulo-Forward 5’ ttgatcgcatggtggctgat 3’ and reverse 5’ gccgtgtcaagtgtttgctt 3’ (124 bp), Lulo-H90-Forward 5’cgtactgggacgtgacaatg 3’ and reverse 5’ gtgacaaaggagggtctgga3’ (184bp), GAPDH-Forward 5’ ttcgcagaagacagtgatgg 3’ and reverse 5’ cccttcatcggtctggacta 3’(150bp). Amplifications occurred under the following conditions: 94 °C / 10min, followed by 40 cycles (94°C / 30sec, 60°C / 30sec, 72°C / 45sec). At the end of the amplification, the dissociation curves in the amplified samples were evaluated to verify possible contaminations or formation of dimers. Q-PCR analysis was performed on the ABI PRISM 7500 Sequence Detection System (Applied Biosystems), and the relative expression of each gene was measured according to the 2-ΔΔCt method (Livak and Schmittgen, 2001) using the constitutive gene GAPDH (GenBank: ACPB02038754) as the internal control of reactions. The gene expression of the heat shock proteins HSP90 and HSP70 in *Lu. longipalpis* was evaluated after the following treatments: females exposed for 1 h at different temperatures (4, 37 and 42°C) and evaluated after 2 h of rest at room temperature (25°C). Females of the same age maintained at 25°C were used as the control group. Groups of females were also evaluated at 0, 2 and 24 hours, and 6 days PIB and PIL.

### 2.9. Statistical analysis

All data were tested for normality using the Shapiro-Wilk test. Multiple comparisons of two or more factors were made with two-way analysis of variance-ANOVA (after normalisation of data when necessary) and Tukey’s multiple comparisons test to confirm where the differences occurred between groups. Multiple comparisons of one factor were evaluated by the Kruskal-Wallis test followed by Uncorrected Dunn’s test. Pairwise comparisons were performed by Mann-Whitney. Differences in proportions among groups were evaluated by the Chi-Square test. In this work results were expressed as the group mean ± the standard error of the mean (SEM), significance was accepted at least when p < 0.05 and * denotes a significant difference. Data was analysed using the software GraphPad Prism (version 7.0, USA).

## 3. Results

### 3.1. Thermopreference

Preliminary trials using 4 day old *-*SF females and males with the thermogradient turned-off showed that distribution of these insects is not dependent on sex (N=40, Two-way ANOVA, n.s). Further, no significance for specific zones of arenas was found in males or females that could suggest thigmotaxis (Tukey’s multiple comparisons test, p> 0.05) **(Figure 2A)**. The activity analysis of these insects showed that females walk further and faster than males (N=40, Mann-Whitney test, p< 0.05), as seen in **Figures 2C, 2D**. When the temperature gradient was established, the distribution of females and males remained similar in the arenas, indicating that the sex of *Lu. longipalpis* did not affect temperature choices of the insect under the influence of a SF diet and such age (N=40, Two-way ANOVA, interaction, n.s). The post hoc test indicated that regardless the sex, *Lu. longipalpis* spend significant time in some specific zones of cold and warm extremities (Tukey’s multiple comparisons test, 1 *vs* 2 and 2, 4, 5 *vs 6*, p <0.05) (**Figure 2B).** Likewise, when the gradient was turned-off, females were more active than males considering velocity and distance moved within arenas (N=40, Mann-Whitney test, p< 0.05) (**Figures 2E** and **2F**).

**Figure 2:**
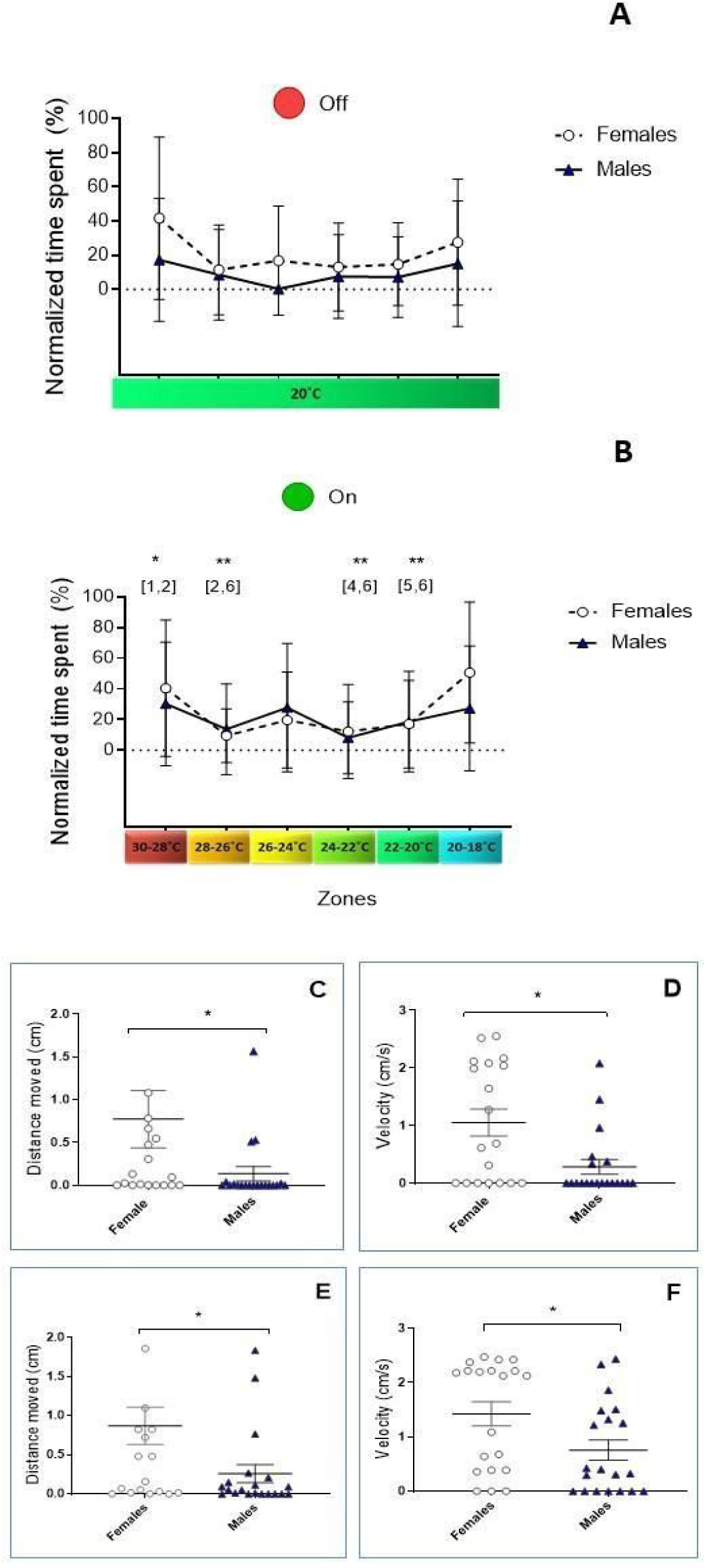
Locomotor behaviour of 4-day-old males and females of *Lu. longipalpis* after feeding with sugar. All insects were maintained in cages under the same temperature (24 °C) before the experiment. The insects were evaluated on a thermal gradient while the apparatus was turned off or turned on. **A** and **B**: Time spent by females and males among the zones; **C**: Average distance travelled by males and females with apparatus turned off; **D**: Mean velocity of movements by males and females with apparatus turned off; **E**: Average distance travelled by males and females with apparatus turned on; **F**: Mean velocity of movements by males and females with apparatus turned on. Range of temperature across zones: 1(30-28°C); 2(28-26°C); 3(26-24°C); 4(24-22°C); 5(22-20°C); 6(20-18°C). Bars represent the difference between groups (Mann Whitney test, * P<0.05) ^a^ at room temperature (20 °C).

The comparison between females 3 days PIL or PIB indicated that *Le. mexicana* did not affect the behaviour of insects at this point of infection (N=40, Two-way ANOVA, interaction, n.s). No differences were found in both groups’ means, and the analysis post hoc demonstrated a preference for all these females for one of the zones of the thermogradient. Blood fed and infected females on the third day spent more time on the cold extremity than other sections of the thermal gradient (Tukey’s multiple comparisons test, 6 *vs* 1,2,3,4,5, p <0.05) (**Figure 3A)**. **Figures 3B** and **3C** display the distance walked and the velocity of movements, both parameters were also similar between infected and non-infected females (N=40, Mann-Whitney test, p< 0.05).

**Figure 3.**
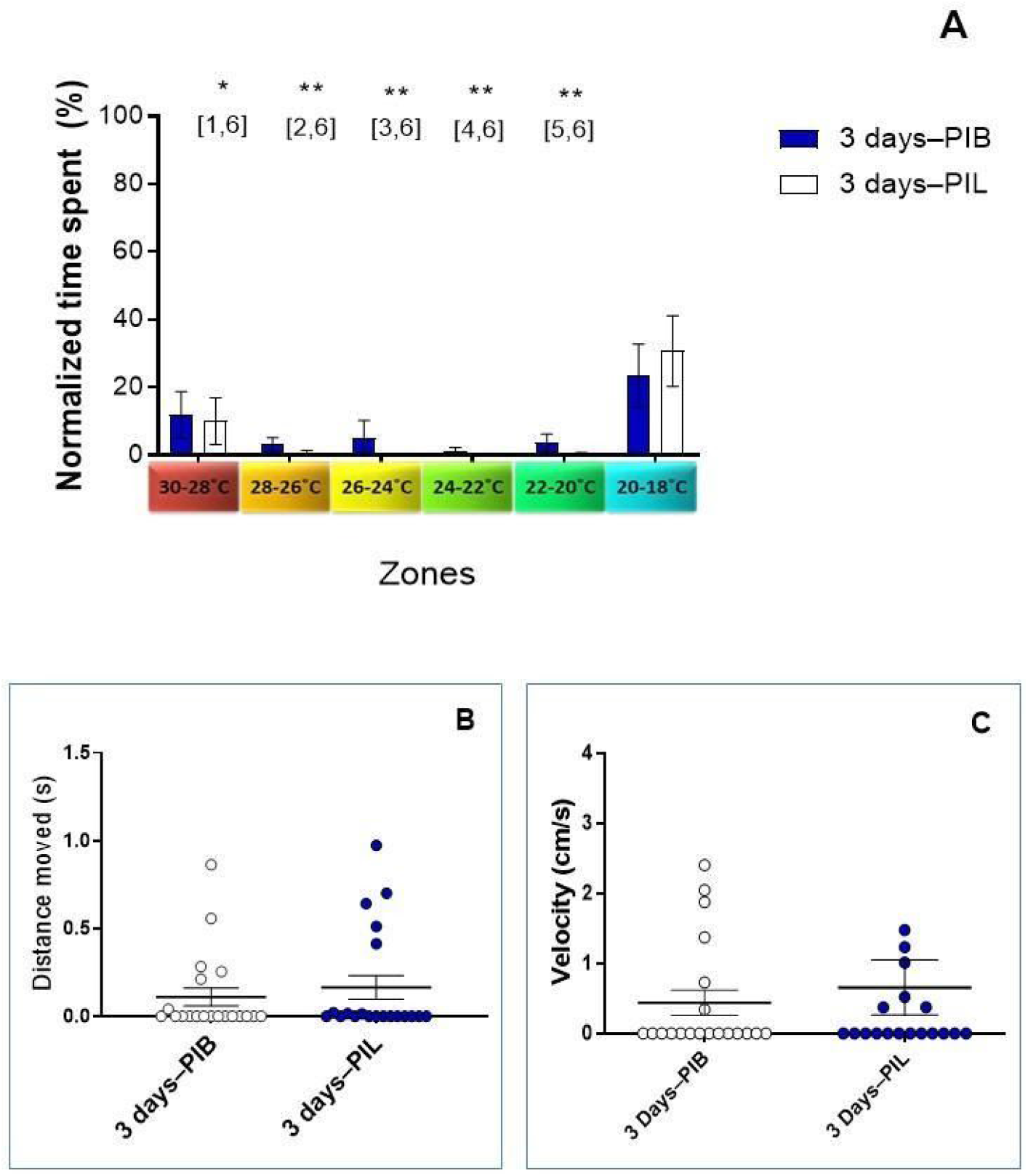
Locomotor behaviour of females on thermal gradient three days after the blood-feeding or infection with *Le. mexicana*. All females were maintained in cages under the same temperature (24 °C) and with free access to sugar before the experiment. The blood or blood + amastigotes of *Le. mexicana* were artificially offered to the females on the 4th day after their hatching. Engorged females from each group were transferred on the same day to different cages, and their behaviour was evaluated in the thermal gradient three days after the procedure. **A** Profile of time that the infected or non-infected females spent among zones. **B** Average distance travelled by females 3 days PIB and PIL. **C** Mean velocity of movements by females 3 days PIB and PIL. PIB: post-ingestion of blood; PIL: post-infection with *Leishmania*. Range of temperature across zones: 1 (30-28 °C); 2 (28-26 °C); 3 (26-24 °C); 4 (24-22 °C); 5 (22-20 °C); 6 (20-18 °C). Numbers in brackets indicate the zone-pair with significant difference (Tukey’s multiple comparisons test), * P<0.05, **P <0.01.

Females were evaluated 6 days-PIB/PIL and again the presence of *Le. mexicana* was not a factor that could affect the preference for temperatures (N=40, Two-way ANOVA, interaction,n.s) **(Figure 4A).** On 6th day PIB/PIL, the behaviour of both groups remained indistinguishable on arenas, but no longer specific to the cold extremity. These females also spent a significant amount of time in the hot extremity of the thermal gradient (Tukey’s multiple comparisons test, 1 *vs* 2, 2 *vs* 5, 3 *vs* 6, 6 *vs* 2, p <0.05), see **Figure 4A.** The activity level was not affected by the infection, females 6 days PIB/PIL walked equally, with the same speed and travel distance as seen in **Figures 4B, C**(N=40, Mann-Whitney test, p>0.05).

**Figure 4.**
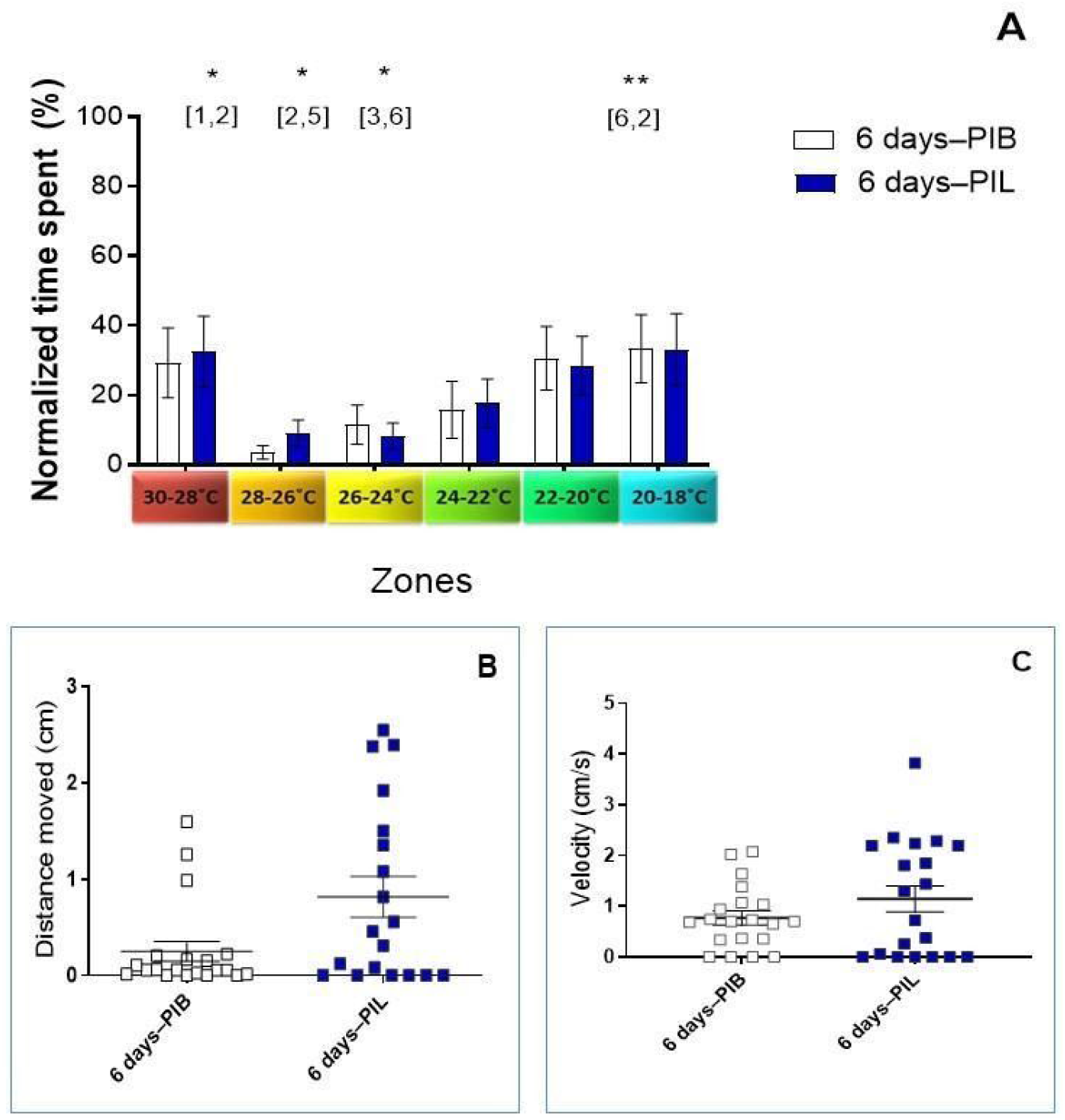
Locomotor behaviour of females on thermal gradient six days after the blood-feeding or infection with *Le. mexicana*. All females were maintained in cages under the same temperature (24 °C) and with free access to sugar before the experiment. Blood or blood + amastigotes of *Le. mexicana* were artificially offered to the females on the 4th day after their hatching. Engorged females from each group were transferred on the same day to different cages, and their behaviour was evaluated in the thermal gradient six days after the procedure. **A** Comparison of time that females infected or non-infected spent among zones. **B** Average distance travelled by females 6 days PIB and PIL. **C** Mean velocity of movements by females 6 days PIB and PIL. PIB: post-ingestion of blood; PIL: post-infection with *Leishmania*. Range of temperature across zones: 1 (30-28 °C); 2 (28-26 °C); 3 (26-24 °C); 4 (24-22 °C); 5 (22-20 °C); 6 (20-18 °C). Numbers in brackets indicate the zone-pair with significant difference (Tukey’s multiple comparisons test) * P<0.05, **P <0.01.

The behaviour of females on the thermal gradient was indifferent under the infection of *Le. mexicana.* Then, we also evaluated data from the same insects considering the time after the procedure as a factor of comparison to find if the age of females could be responsible for their choices. However, no interference of the age in the PIL group was found that could support this hypothesis (N=40, Two-way ANOVA, interaction, n.s). Tukey post hoc tests revealed that mean values are different between females 3-days PIL and 6-days PIL when the zone distribution is ignored (p<0.05). All these infected females, independent of age, spent more time and had a higher frequency in the cold area of the thermogradient (Tukey’s multiple comparisons test, 6 *vs* 2, 3, 4 p <0.05), **Figure 5A.** A meaningful contrast between both groups was seen looking at the activity, since infected *Lu. longipalpis* females were slower and walked less on the third day PIL (Mann-Whitney test, p<0.05) as shown in **Figure 5B, C.**

**Figure 5.**
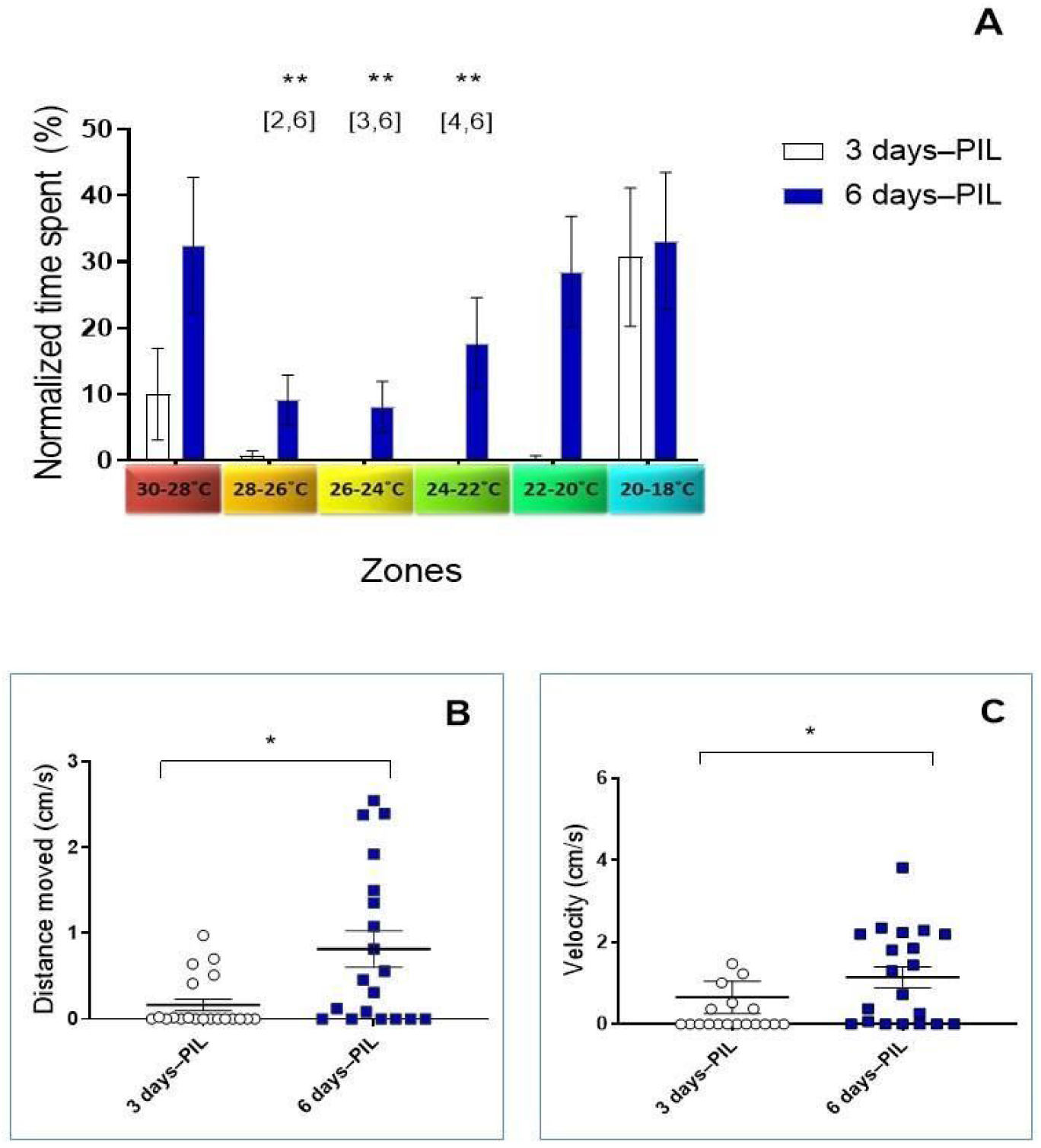
Locomotor behaviour of females on thermal gradient three and six days after infection with *Le. mexicana*. All females were maintained in cages under the same temperature (24 °C) and with free access to sugar before the experiment. Blood with *Le. mexicana* was artificially offered to the females on the 4th day after their hatching. Engorged females from each group were transferred on the same day to different cages, and their behaviour was evaluated in the thermal gradient on third and sixth days after the procedure. **A** Comparison of time spent among zones between females 3 and 6 days old PIL. **B** Average distance travelled by females 3 and 6 days PIL. **C** Mean velocity of movements by females 3 and 6 days PIL. PIL: post-infection with *Leishmania*. Range of temperature across zones: 1 (30-28 °C); 2 (28-26 °C); 3 (26-24 °C); 4 (24-22 °C); 5 (22-20 °C); 6 (20-18 °C). Numbers in brackets indicate zones pairs with significant difference (Tukey’s multiple comparisons test) and bars represent the difference between groups (Mann Whitney test), * P<0.05, ** P <0.01.

Similarly, the age of females after a blood meal have not affected choices of temperatures (N=40, Two-way ANOVA, interaction, n.s). Means values between females 3 and 6 days after blood feeding also were distinct (p<0.05) (**Figure 6A**). Regardless of age, blood-fed *Lu. longipalpis* spent more time and visited more frequently the cold area of arenas (Tukey’s multiple comparisons test, 6 *vs* 2,3,4 p <0.05). The activity of these females also was similar to the profile found in infected females. On 3rd-day the females were slower and walked less than on the 6th-day after the feeding (Mann-Whitney test, p<0.05) (**Figure 6B, C**)

**Figure 6.**
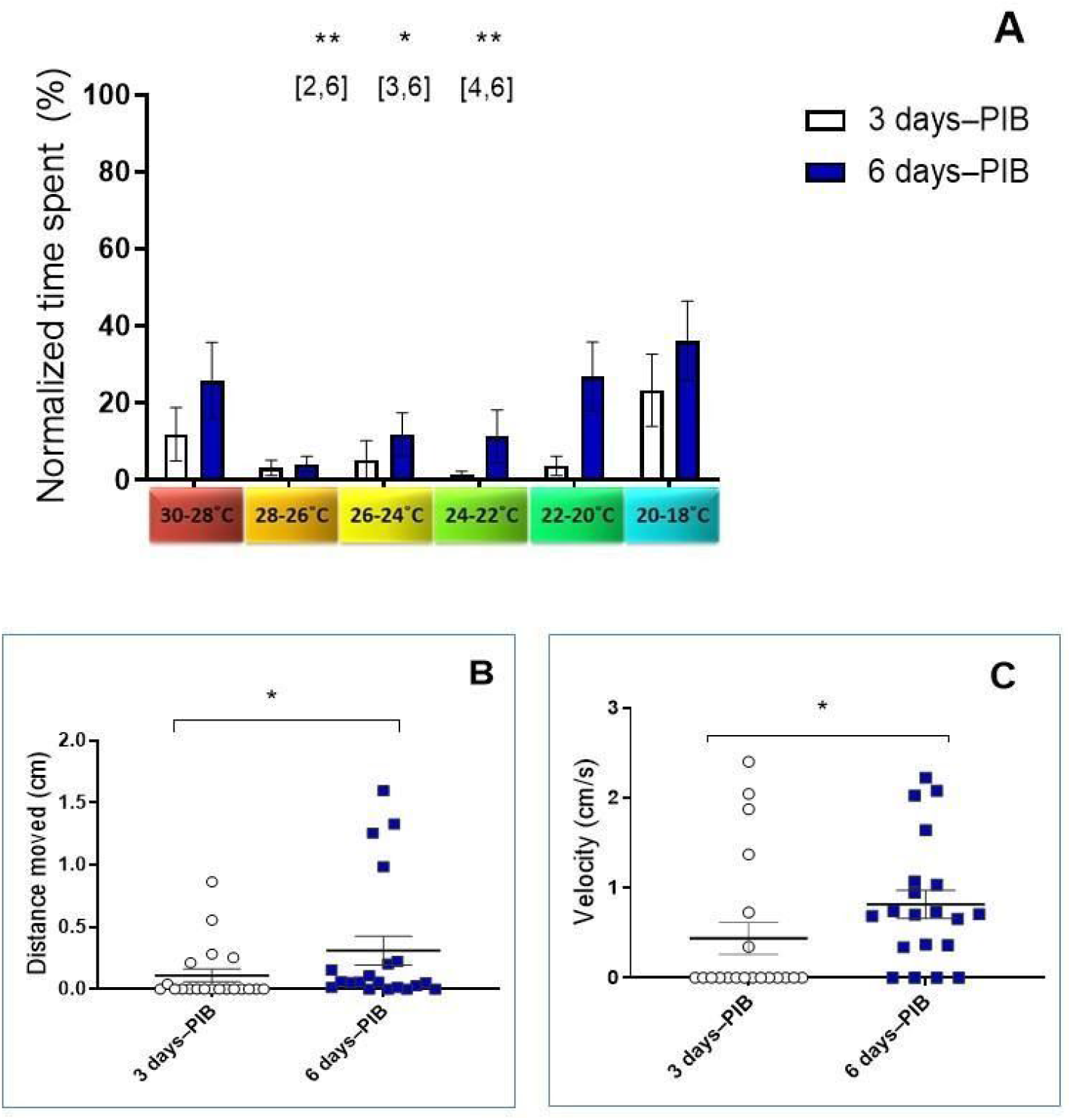
Locomotor behaviour of females on thermal gradient three and six days after the blood-feeding. All females were maintained in cages under the same temperature (24°C) and with free access to sugar before the experiment. Blood was artificially offered to the females on the 4th day after their hatching. Engorged females from each group were transferred on the same day to different cages, and their behaviour was evaluated in the thermal gradient on 3rd and 6th days after the procedure. **A** Comparison of time that females spent among zones. **B** Average distance travelled by females 3 and 6 days PIB. **C** Mean velocity of movements by females 3 and 6 days PIB. PIB: post-ingestion of blood. Range of temperature across zones: 1 (30-28°C); 2 (28-26°C); 3 (26-24°C); 4 (24-22°C); 5 (22-20°C); 6 (20-18°C). Numbers in brackets indicate zones pairs with significant difference (Tukey’s multiple comparisons test) and bars represent the difference between groups (Mann Whitney test), * P<0.05, **P <0.01.

The contrast between females on the 3rd and 6th day after the blood ingestion continued when the post-oviposition of females was followed. Females that were kept at the same temperature as those used on the thermal gradient (24°C) were able to lay eggs only in 10% of pots on the third day after the blood-feeding. In contrast, on the sixth day, the proportion increased significantly to 75% considering the total number of pots (N=20, Chi-square test, p<0.01). The temperature influenced both the oviposition and mortality proportions of *Lu. longipalpis* females, as seen in **Table 1**. On the third day after blood feeding, females carried at 30 °C had no remains of blood inside their gut, oocytes were in the advanced level of development, and all females were found dead, leaving eggs in 85% of pots (**Supplementary Figure 2**). In contrast, females’ three days PIB at 20°C remained with a large amount of undigested blood inside the gut and oocytes were still poorly developed. First eggs inside these pots were seen only 4 days PIB, but the survival of females 6 days after oviposition was higher compared with those carried at 24°C or 30°C (N=60, Chi-square test, P <0.05).

**Table 1.** Effect of temperatures on oviposition and mortality of females of *Lu. longipalpis* after the blood-feeding. The presence of eggs and mortality was followed until the 6th day after the blood-feeding.

| Day after the blood feeding |  | Temperatures N (%) |  |  | Chi-square, df | P value |
| --- | --- | --- | --- | --- | --- | --- |
|  |  | 20°C | 24°C | 30°C |  |  |
| Number of eggs | 3rd | 0 (0) | 2 (10) | 17 (85) | 9.604, 2 | 0,0082* |
|  | 6th | 4 (20) | 15(75) | 17 (85) |  |  |
| Mortality | 3rd | 1(5) | 2 (10) | 20 (100) | 6.182, 2 | 0,0456* |
|  | 6th | 2 (10) | 13 (65) | 20 (100) |  |  |

As expected, temperatures also affected the development of *Le. mexicana.* First flagellate forms of parasites were found just on 4th day after infection for the sand flies maintained at 20°C, and 3 days for those kept at 24°C. Promastigotes were not detected in females kept at 30°C. The proportion of infected females after 6 days of infection changed according to temperature; 20°C (85%), 24°C (70%), 30°C (0%), n=30.

### 3.2. Thermography

After some probing bites, females of *Lu. longipalpis* started to ingest the blood from mice remaining in the same place until the end of the feeding. During this period, it was possible to see an increase in the body temperature of insects a few seconds after the landing. This increase occurred homogeneously, and then the temperature remained constant around 35 ± 0.5 °C during all blood ingestion (n=10), **Figure 7C**. In the final moments of feeding, when there was urine release, we also did not observe any changes in body temperature of females (**Figure 7D**).

**Figure 7.**
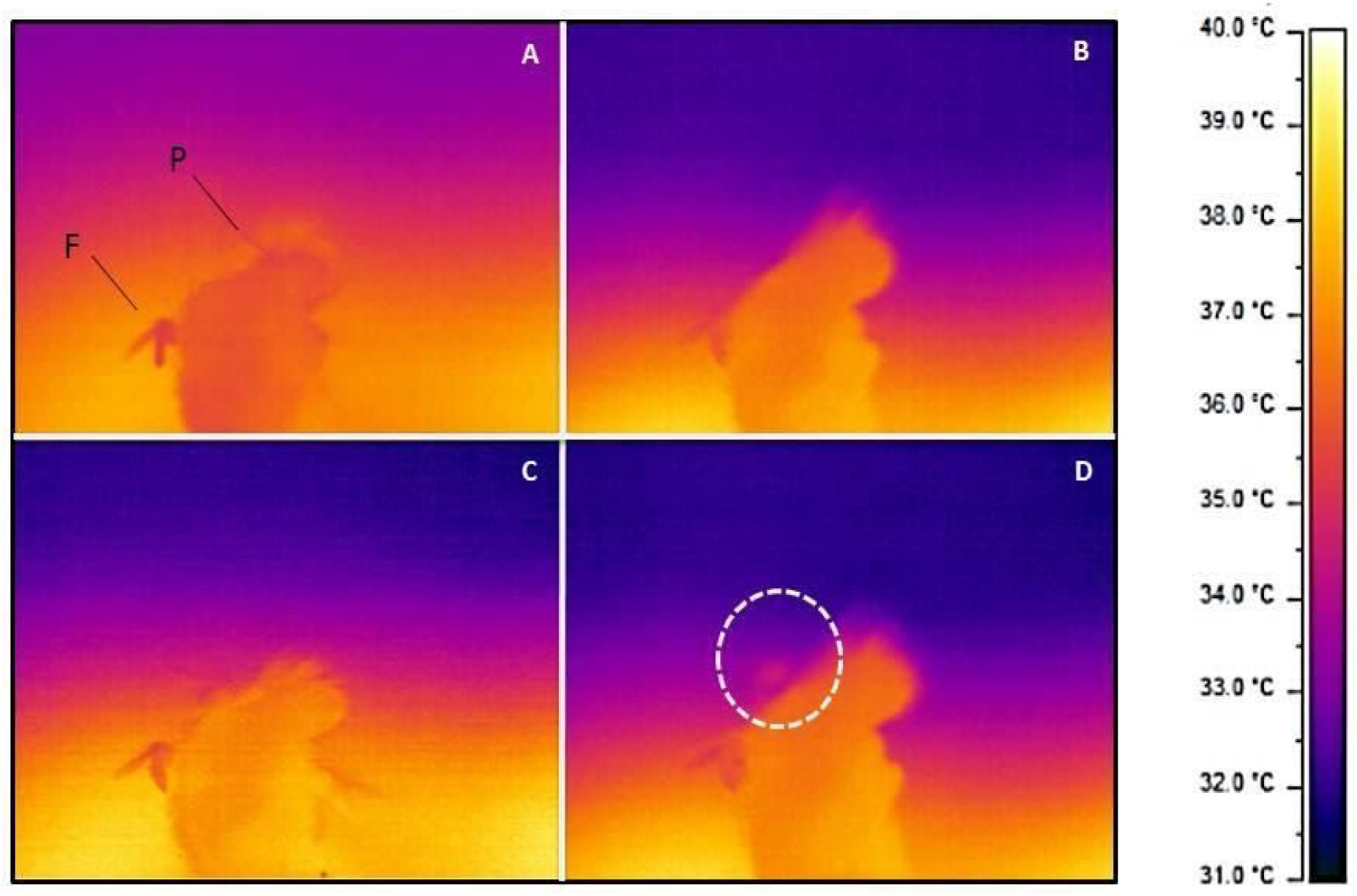
Thermography of the *Lu. longipalpis* female during blood-feeding on a mammal’s host. **A** Female just after landing on the skin (before bite). Here, P denotes the paw of the mouse, F denotes the female of *Lu. longipalpis*. **B** Initial feeding moment (after the bite). **C** Female showing abdomen already distended by blood ingestion. **D** Female at the final moments of feeding with the release of urine. Dashed circle highlights a droplet of urine released in the opposite direction of feeding.

### 3.3. Heat shock proteins

The exposition of adult females to temperatures tested (4 °C, 37 °C, 40 °C) induced a distinct expression of HSP90(83) (Kruskal-Wallis test, p<0.05). The stress caused by heat at 40 °C stimulated an increase of 4-fold in HSP90 expression compared with those remaining at room temperature (Dunn’s test, p < 0.05), **Figure 8A**. However, no differences were found when the same temperatures and insects were tested for HSP70 transcripts (Kruskal-Wallis test, p>0.05), Figure 8B.

**Figure 8.**
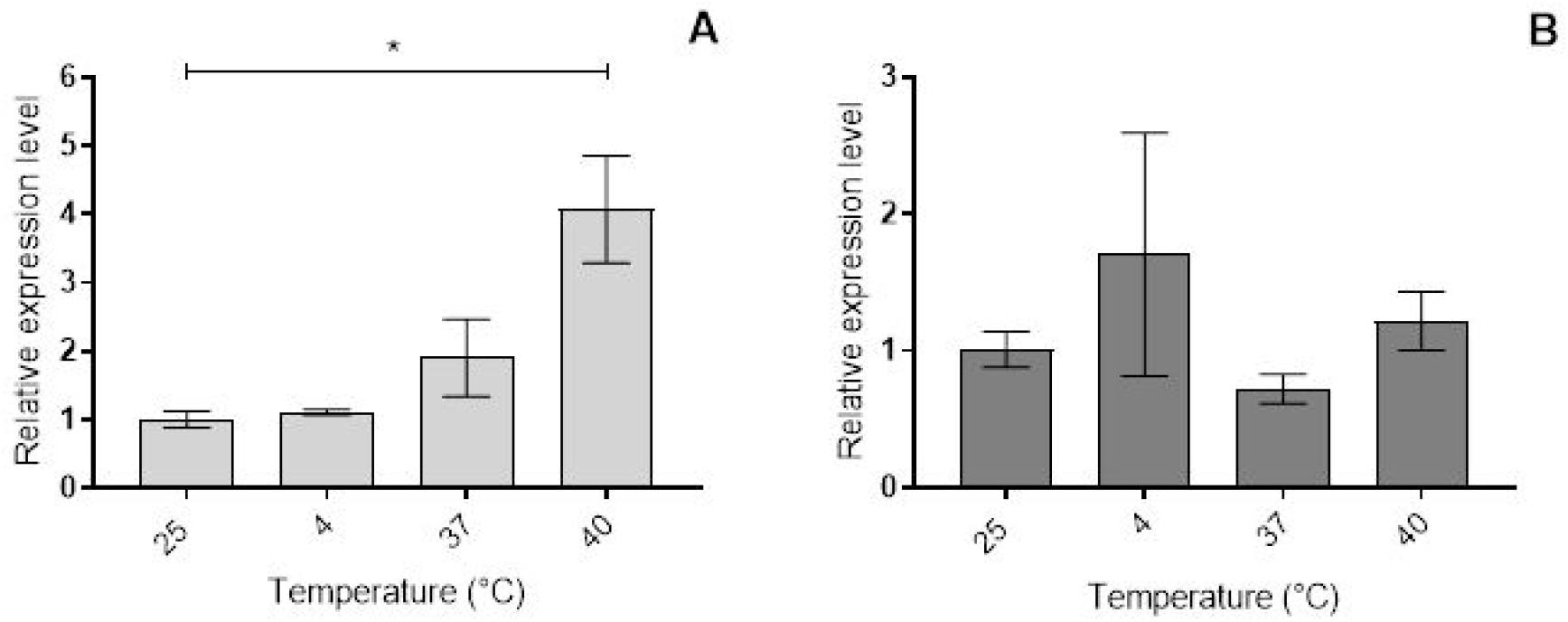
Levels of heat shock proteins HSP90(83) and HSP70 in females of *Lu. longipalpis* exposed to different temperatures. Females sugar-feeding were exposed to different temperatures for 1 hour and left to rest for 2 hours. **A** The relative mRNA expression level of the hsp90. **B** The relative mRNA expression level of the hsp70. Bar charts represent the mean ± SEM of three pools of 10 females collected during three different temperature exposition events (Kruskal-Wallis test, p<0.05).

Levels of the same heat shock proteins were tested in females blood-fed and infected with *Le. mexicana,* to evaluate whether the stress caused by the parasite could affect the expression of HSP-coding genes. Comparisons between PIB and PIL showed a significant difference of profile when looking at the expression of HSP90(83) (Two-way ANOVA, interaction, P<0.05). Females-PIB decreased levels of expression faster when compared to females-PIL, as seen in **Figure 9A.** The same analysis was performed looking for HSP70 gene expression, but no differences were found considering the procedure and time frame (Two-way ANOVA, interaction, n.s).

**Figure 9.**
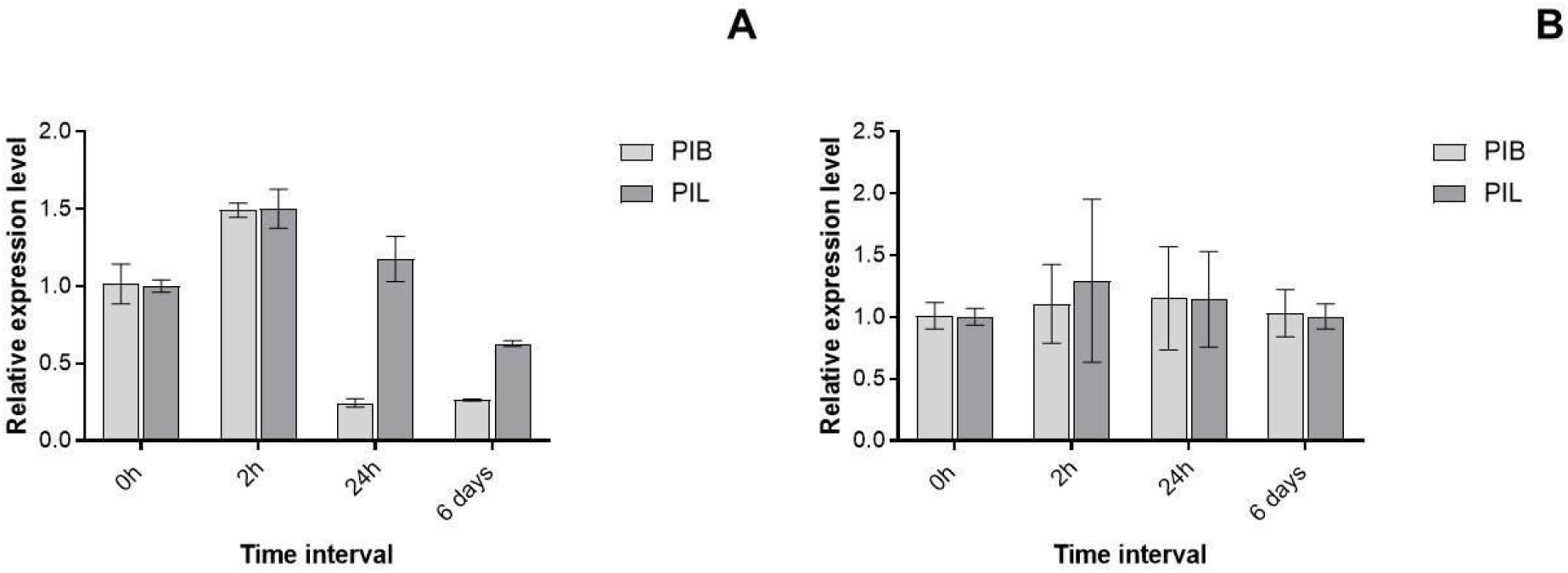
Profile of relative transcripts levels of heat shock proteins HSP90(83) (**A**) and HSP70 (**B**) in females of *Lu. longipalpis* after blood ingestion or infection with *Le. mexicana*. PIB: post-ingestion of blood; PIL: post-infection with *Leishmania*. Bar charts represent the mean ± SEM of three pools of 10 females collected during three different blood ingestions/infections (Two-way ANOVA, P<0.05). The average Δct at time 0h from each group was used as the reference value to normalise all subsequent time points.

## 4. Discussion

In this work, we explored the thermal sensibility of the sand fly *Lu. longipalpis* in different physiological conditions. We found that sand flies 4-days-old did not manifest preference for specific areas of arenas without temperature stimulus. However, insects in the same conditions started to spend more time in cold and warm zones when the temperature gradient was stabilised. The activity evaluated in these groups showed that females were more active than males; such sexual dimorphism has also been observed in other insects as Drosophila (Martin, 2004). Three days after blood ingestion or infection, females spend more time in the coldest zone (20-18°C), besides to become less active. On the other hand, on sixth-day females in the same conditions, also started to spend more intervals at the warmer extremity. Besides that, females on day 6 were more active, walking more and faster.

Although the day of procedure had no impact on temperature choices, there is a clear distinction between a group of females on day 3 and 6 after the procedure, independent of zones. Reduced mean values across arenas were obtained from females three days after procedures, and it is probably a repercussion of the lethargy found in females from this group **(Figures 5 and 6).**

An additional comparison between females 3 days PIB and sugar-fed females with same age (7 days old) confirmed that the sluggish behaviour of females 3 days PIB was not related to age, but possibly to the diet of females (N=40, Two-way ANOVA, interaction p<0.05). Furthermore, the activity of sugar-fed females is higher than blood-fed females with the same age (Distance moved/ velocity, Mann-Whitney test, p<0.05), **Supplementary Figure 1.**

The distinction between females on days 3 and 6 after blood ingestion were also found through observations of the oviposition rate of females carried at the same conditions as those used on thermal gradients. While on day three, only 10% of females had laid eggs, on the sixth-day majority of pots already had eggs **(Table 1).** Also, dissections of some females after experiments on thermogradient confirmed the contrast between females visually from both days. On day 3 PIB/PIL it was possible to see a higher number of developed oocytes compared with those dissected on day 6. These observations are supported by previous work observing females of *Lu. longipalpis* maintained at 25 °C. Magnarelli (1984) showed that only 32% of *Lu. longipalpis* contained mature oocytes three days after a blood meal, and by the 6th day the proportion increased to 69%. It suggests that the weak activity and preferences of 3 days PIB/PIL for the cold zone can be related to its oviposition condition.

Insects tend to look for oviposition places that will support their further offspring. However, previous experiments performed by Hilda and Tesh (2000) suggest that such range of 20-18°C chosen by 3 days PIB/PIL is not optimal for oviposition, egg’s hatching, the larval development and adults’ emergence of *Lu. longipalpis*. In contrast, the same work found an increase of longevity for the species in temperatures even lower (15°C). Considering that environmental conditions offered on the surface of arenas did not favour the oviposition of sand fly females, it is possible that behaviour of gravid females 3 days PIB or PIL was linked to their self-preservation until to find an appropriate site to oviposition. As many other sand flies, *Lu. longipalpis* are found in tropical areas and their putative breeding sites are usually unseen. When found in nature, immature forms of this vector are generally related to hidden places: under rocks, at the base of trees, under leaves, rodent burrows and animal shelters (Deane and Deane, 1957; Ferro et al., 1997). Ectotherms are incapable of surviving in open habitats through physiological thermal tolerance alone, and thus they must have access to thermal shelter to survive (Sundaya et al., 2014). Microenvironments like that should also be essential to gravid females to prevent their bodies overheating, especially considering their fragile condition. This hypothesis is supported especially by the slow velocity and shorts distance moved by females 3 days after PIB/PIL. This activity results are compatible with findings of Meireles-Filho et al. (2006). In this work, the locomotor activity of *Lu. longipalpis* in the first 48 hours after blood feeding was reduced around 40% when compared to non-fed females.

Unlike females on the third day, most females on the sixth day had already laid their eggs, and previous work from our group showed that it is an essential condition to females perform the next blood feed (Moraes et al., 2018). Therefore, the higher levels of activity and additional orientation for the hottest section of the arena may reflect the release of eggs and the search for a new source of blood by these females on the sixth day. These contrasting choices of cold and hot sections (**Figure 4)** may reflect the probing behaviour of females on their vertebrate host. But it is essential to consider that the behaviour of all groups was analysed since the first 5 minutes after placed on arenas. Some initial trials using 4 days old sugar feeding insects on turned off apparatus showed that after exploring areas, *Lu. longipalpis* settled down only after ±15min. Therefore, data analysed on thermal gradient are representing not exclusively the final resting position selected by insects, but also the exploratory behaviour. On the other hand, the shorter and slower distance moved by groups like 3 days PIL/PIB females since the first minutes suggests that exploratory behaviour can change depending on the physiological condition of the insect. For vulnerable females 3 days PIL/PIB, saving energy is probably more critical than exploring the environment for extended periods. Therefore, the influence of the exploring period on analysis of time spent in zones can be reduced in this group, while is more intense in groups of high activity as females 6 days PIL/PIB. This method used for live tracking could be a drawback to evaluate the resting position of insects. However, other important aspects of sand fly behaviour would be lost observing only the final resting position, like the activity behaviour among groups.

The similar activity and thermal sensibility of females 6 PIB/PIL suggest that *Le. mexicana* has not triggered the behavioural immunity responses of the insect host. Even so, the higher activity of all females at day 6 compared to day 3 also means that infected females will be more active exactly when parasites are found in the stomodaeal valve. Therefore, the higher chances of transmission of the parasite at this point of infection are supported by the active behaviour of the insect in response to its stage of blood digestion. Although *Lu. longipalpis* is not a natural vector of *Le. mexicana*, this species of *Leishmania* is a well-known model for infection in this insect. Moreover, when infections are carried at similar temperatures, *Le. mexicana* produces a larger density of parasites in the gut when compared to *Le. infantum* on the sixth day of infection (Moraes, 2018; Serafim, 2018). Therefore, we believe that if *L. infantum* could trigger the temperature-related behaviour of *L. longipalpis* we would observe such alterations during our trials. The results reported here are similar to those seen in *Anopheles stephensi*, in which the insect was able to thermoregulate behaviourally, but could not respond to infections with fungal entomopathogens or *Plasmodium yoelii* (Blanford, 2009).

During the behaviour experiments on the thermogradient, all females were maintained in cages at 24 °C before experiments. Since the temperature affects the speed of blood digestion, it is also important to consider that females maintained at different temperatures could generate a distinct behaviour. Our results showed that mild or warmer temperatures could retard or accelerate female mortality and oviposition. This effect also can be seen visually after dissecting some females maintained at 24 °C, 30 °C and 20 °C **(Supplementary Figure 2).** Moreover, this becomes especially important among infected females since parasite development and insect survival were also directly affected by temperature. To clarify, females kept at 20°C on the day 6 post-infection would be holding a large amount of retained eggs, which could result in females being less avid and sensible to the host temperature. In addition, these females in this same day would probably have a reduced chance supporting parasites in metacyclic forms due to the slower development. Thus, some extra days would be needed for these females to fulfil their role as a vector and to transmit the parasite.

To perform their feeding, many phytophagous insects avoid heat from solar radiation by changing their behaviour (Hochuli, 2001; Willmer, 1983). However, in the case of hematophagous insects it is more challenging for them to change behaviour while performing the blood-feeding. Therefore, the ingestion of blood must be performed by an insect in the most efficient way (Lehane, 1991). If the feeding is interrupted by intense locomotor activity in an attempt to escape the heat from the host, then this behaviour may increase the exposition time of an insect. As a result, the risk of not having a whole blood meal, or even being killed by the vertebrate host will increase. Lahondère & Lazzari (2012) and Lahondère *et al.* (2017) have described that some blood-sucking insects possess morpho-physiological adaptations allowing them to avoid overheating when they feed on a warm-blooded vertebrate. For instance, while ingesting blood the triatomine *Rhodnius prolixus* uses a sophisticated heat exchanger inside its head to dissipate the excess of heat and to maintain the abdominal temperature close to the ambient temperature. In the case of *Anopheles* mosquitoes, the decrease of the abdominal temperature happens by evaporation of excreta released during blood feeding. Observations of thermography in *Lu. longipalpis* indicated that in a short time after the start of ingestion of the blood the body temperature of females reached the same temperature as that of the host. Moreover, the temperature was homogeneous in the whole body of the insect, and no thermal related behavioural mechanisms were observed during feeding. Thus, it may be important for the physiology of *Lu. longipalpis* to rely on other mechanisms of adaptation, such as the expression of heat shock proteins. In our results females of *Lu. longipalpis* showed an increase of HSP90(83) expression, being significant when females were exposed only to 40 °C. However, the expression level increased 1-fold at 37 °C, suggesting that levels of expression can be temperature-dependent **(Figure 8)**. After the blood-feeding or infection at 37°C there was also a subtle increase of expression of the same gene, with a peak 2h hours after the procedure. This level is consistent when females were exposed at 37 and evaluated 2 hours later after the procedure. A difference between infected and non-infected females was also observed at 24hours, however, this difference is related to a sharp decrease of gene expression in blood-fed females while compared to the subtle reduction in the group of infected females.

The increased expression of heat shock proteins due to the ingestion of warm blood has been studied in other vectors. It was found that the knockdown of HSP70 generated mortality in mosquitoes and kissing bugs after the ingestion of the blood (Paim et al., 2016; Benoit et al., 2011). The level of this protein (HSP70) was not significant during our experiments, but the expression of these chaperones in relation to temperature stress can be extremely variable from one species to another (Zhao and Jones, 2012). In this work we explored for the first time the expression of HSP’s in *Lu. longipalpis*. Our results suggested that these proteins, potentially HSP90 (83), are relevant for sand flies under temperature stress. However, to confirm this suggestion, it is essential to perform further studies based on probe analysis, such as Northern blots combined with gene knockout to ensure the translation and phenotype of HSP’s.

In nature, sand flies usually have a nocturnal pattern of behaviour and a higher level of activity during evenings (Souza et al., 2005; Feliciangeli et al., 2014; Morrison et al., 1995). This habit should be adaptive, considering pieces of evidence of the high sensitivity of these insects to warmer temperatures. Thus, probably sand flies should control the body temperature by changing their behaviour, i.e. by remaining in shaded and hidden places to avoid the solar radiation and leaving those places only for some essential activities, such as feeding. However, during these moments, the behaviour is probably not the most efficient strategy to control the body temperature of these insects.

## Conclusion

In this work we found that females of sand fly *Lu. longipalpis* adjust their behaviour and body temperature in response to their gonotrophic cycle, and the results suggested a limited effect of *Le. mexicana* in the insect host. In certain moments of thermal stress when thermal behaviour is not used, other physiological responses can be triggered by the insect, as the expression of certain heat shock proteins.

## Acknowledgements

This work has the support of Research Agencies (CNPq/CAPES) linked to Program Science without Borders (Brazil) and from Lancaster University (UK). The authors are grateful for the technical assistance of Michelle Bates.

## Supplementary Figures

**Supplementary Figure 1.**
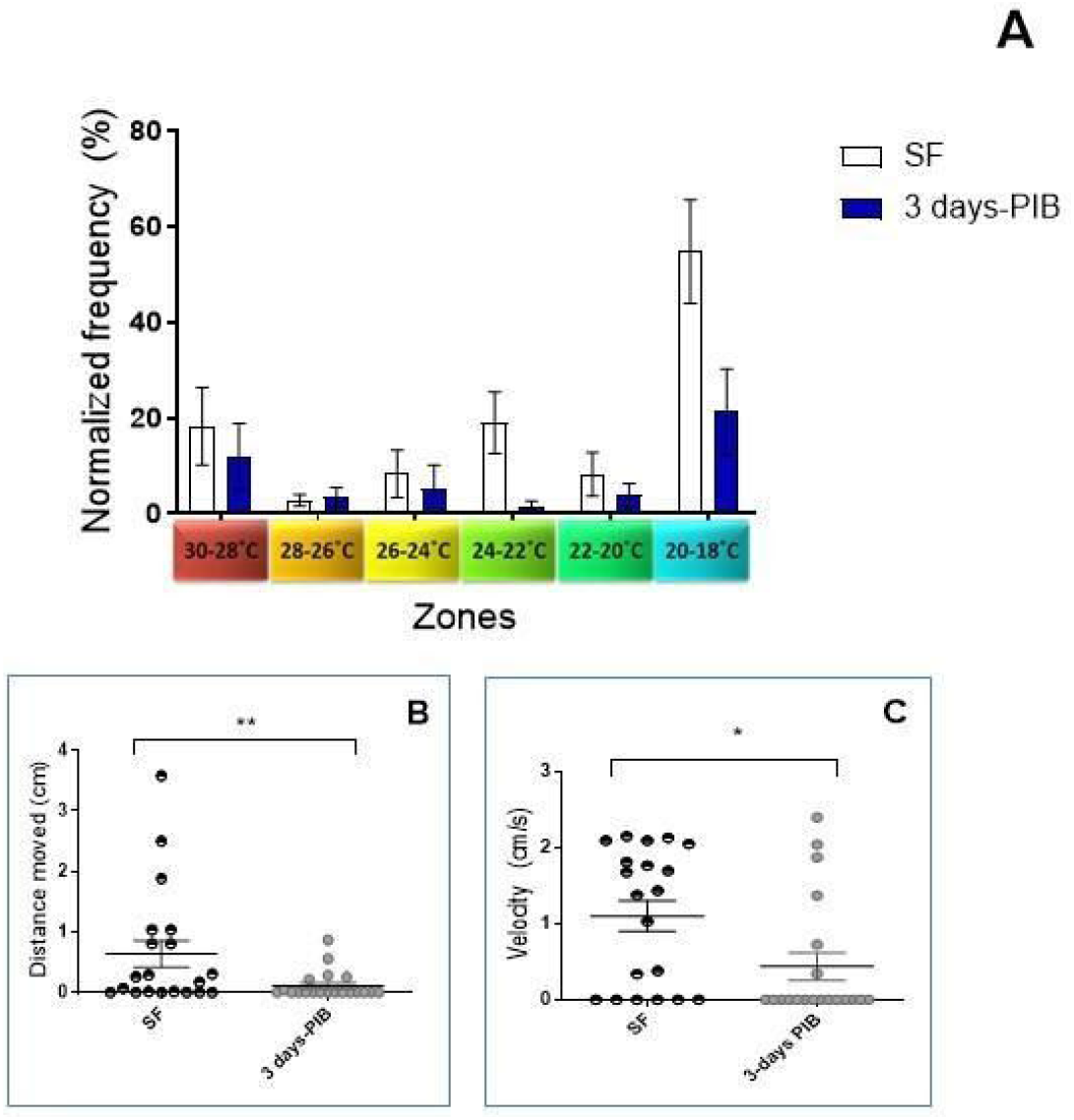
The behaviour of females seven days old on thermal gradient after ingestion of blood or sugar. All females were maintained in cages under the same temperature (24 °C) and with free access to sugar before the experiment. On the 4th day after the hatching, a group of females was fed with blood, and their behaviour was evaluated on thermal gradient three days after the procedure (3 days PIB). On the same day, the behaviour of females maintained only with sugar was evaluated (SF-Sugar feeding). **A** Comparison of the frequency of females spent among zones. **B** Average distance travelled between females SF and 3 days PIB. **C** Mean velocity of movements between females SF and 3 days PIB. Range of temperature across zones: 1 (30-28 °C); 2 (28-26 °C); 3 (26-24 °C); 4 (24-22 °C); 5 (22-20 °C); 6 (20-18 °C).

**Supplementary Figure 2.**
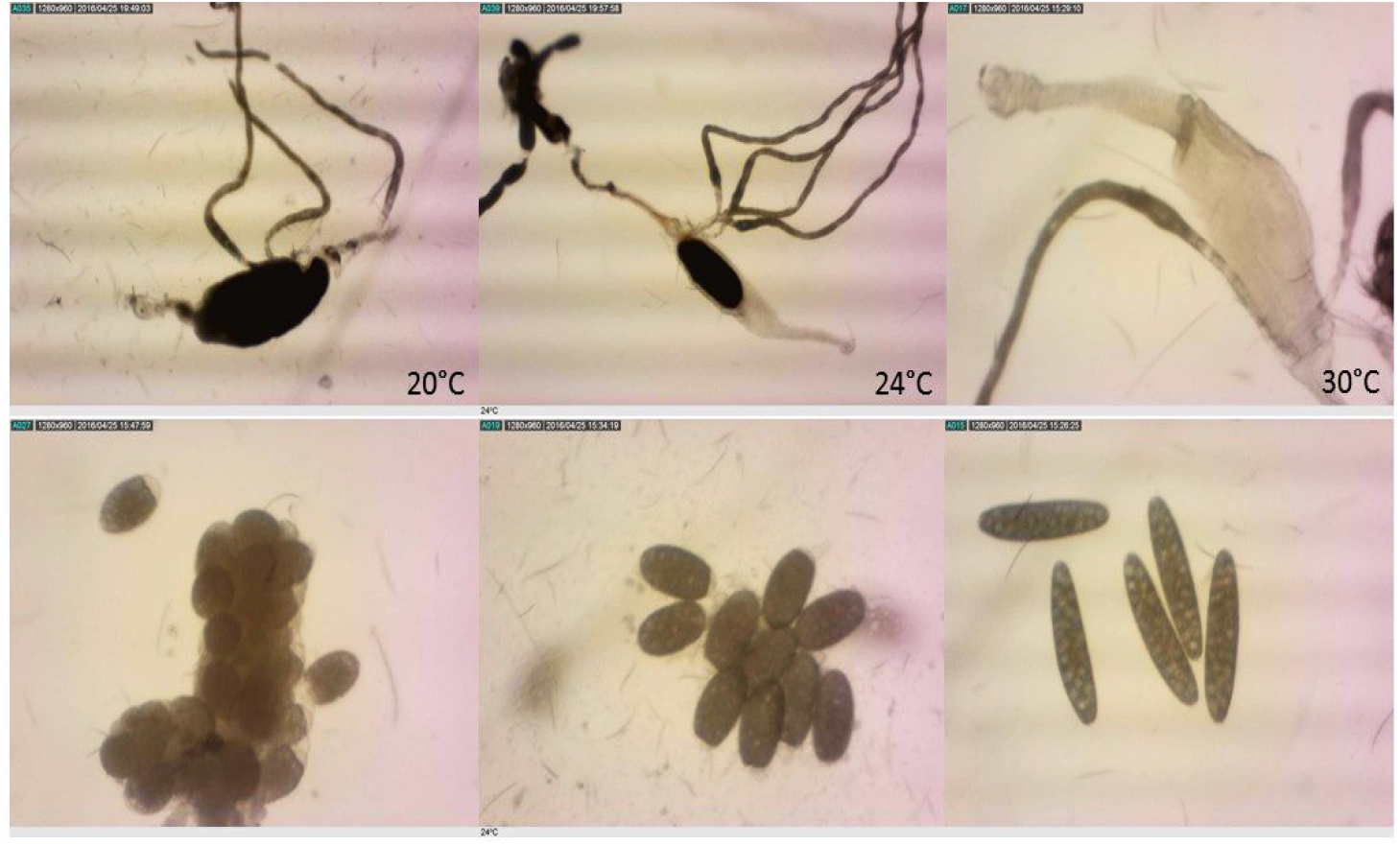
Evolution of blood digestion and eggs development according to temperature in which *Lu. longipalpis* females were maintained. The figure illustrates the gut and oocytes of females 3 days after blood ingestion at 20 °C, 24 °C and 30 °C.

